# MHCXGraph: A Graph-Based approach to detecting T cell receptor cross-reactivity

**DOI:** 10.64898/2026.04.07.717034

**Authors:** Carlos Daniel Marques Santos Simões, Rocio Lucia Beatriz Riveros Maidana, Samuel Chagas de Assis, João V. S. Guerra, Helder Veras Ribeiro-Filho

## Abstract

The T cell receptor (TCR) recognition of multiple peptides presented by the major histocompatibility complex (MHC) is a key natural phenomenon, enabling the T cell repertoire to respond to a broad array of antigens. Despite its importance to the immune response, T cell cross-reactivity poses a major challenge for the development of novel T cell–based therapies. In this study, we present `MHCXGraph`, a graph-based computational approach for identifying conserved and immunologically relevant regions across multiple structures of peptides bound to MHC molecules (pMHC). Our approach provides three operational modes with user-defined parameters, allowing flexible configuration according to specific scientific needs while delivering fully interpretable results through user-friendly interfaces. We evaluated `MHCXGraph` across three case studies, including peptides bound to classical MHC Class I, MHC Class II, and unbound HLA alleles, demonstrating its ability to capture conserved structural determinants beyond sequence similarity. By integrating structural information with efficient graph-based analysis, `MHCXGraph` addresses key limitations of sequence-based methods while maintaining computational scalability. Collectively, these results indicate that `MHCXGraph` can be readily integrated into computational pipelines for T cell cross-reactivity discovery, especially in the context of *de novo* pMHC engager design and T cell–based vaccine development.

## 1 Introduction

T cells, through their surface-expressed T cell receptors (TCRs), recognize short peptides presented by the major histocompatibility complex (MHC) class I or class II molecules (hereafter referred to as pMHC) [1, 2]. The TCR binding to pMHC can trigger T cell activation and constitutes one of the most critical events at the immunological synapse, initiating the adaptive immune response against pathogens and tumor-related antigens. Unlike antibodies, which undergo affinity maturation to enhance their specificity toward target antigens, TCRs exist as a diverse repertoire shaped in the thymus through positive and negative selection [3, 4]. T cell maturation process ensures TCR tolerance to self-peptides while enabling recognition of wide range of non-self peptides, including those derived from viral proteins or tumor-associated mutated proteins (i.e. neoantigens) [5, 6].

Although vital for immune surveillance, the ability of TCRs to recognize multiple peptides presented by MHC, a phenomenon known as cross-reactivity, often limits the therapeutic use of T cells, particularly those engineered with modified TCRs [7, 8]. Engineered TCRs may inadvertently recognize self-peptides in addition to their intended targets, leading to severe adverse effects. A well-known example is a TCR designed to target the melanoma-associated antigen MAGE-A3, which cross-reacted with a peptide derived from the cardiac protein Titin, leading to fatal cardiotoxicity in patients [8]. More recently, Artificial intelligence (AI)-based approaches have enabled the *de novo* design of novel TCR-like binders to pMHC targets, further increasing the need for robust strategies to assess and mitigate cross-reactivity [9, 10]. These challenges also extend to vaccine development, where epitope similarity to self-peptides may lead to immune tolerance and reduced immunogenicity [11].

To address these issues, computational methods have been developed to identify and rapidly assess the potential for cross-reactivity, generally based on the assumption that shared patterns among pMHC complexes may drive recognition by the same TCR. Conventional approaches rely on sequence-based comparisons, in which a target peptide is evaluated against self-peptides derived from the human proteome and ranked according to sequence identity, biochemical properties, and immunopeptidomic data [12]. For instance, ICrossR performs position-wide comparisons of peptide sequences to assess identity [13], whereas sCRAP incorporates physicochemical and structure-derived information, excluding MHC anchor residues that are not accessible for TCR interaction [14]. CrossDome further refines this approach by applying biochemical profiles and weighting peptide positions that may be critical for TCR recognition [15].

Although sequence-based approaches focused on the peptide provide a rapid initial screening, relying solely on peptide sequence information has important limitations. First, the MHC polymorphism, which can strongly influence TCR specificity, is completely overlooked. In addition, while peptide binding to MHC involves conserved pockets that restrict variability in binding modes, shifts in peptide register can occur [16, 17], as well as structural adaptability in peptide conformation [18]. These factors are not captured by current sequence-based comparisons, especially when peptides of different lengths are involved. Thus, to achieve a more accurate comparison between pMHCs, 3D structural information can be employed, which is critical and urgently needed for the robust identification of potential cross-reactivity. This has become even more demanding given recent advances in accurate and high-throughput 3D protein structure prediction with AlphaFold and related structural prediction methods [19]. MatchTope [20] represents one such effort, focusing on pMHC class I and comparing electrostatic surface properties after structural alignment. However, a general, alignment-free framework capable of identifying physicochemical similarities in a structurally interpretable manner, without topological constraints, across diverse pMHC structures remains lacking.

Here, we introduce `MHCXGraph`, a versatile and interpretable computational tool based on graph-theoretical representations of pMHC 3D structures to perform the structural characterization and detection of potential TCR cross-reactivity derived from pMHC surface similarities. `MHCXGraph` is designed to be integrated into cross-reactivity prediction pipelines and extend existing sequence-based approaches by enabling structure-driven analysis. `MHCXGraph` employs an alignment-free strategy based on maximum common subgraph detection, using geometric association graphs to approximate all possible spatial correspondences between protein structures[21, 22, 23]. This approach enables the identification of regions on the pMHC exposed surface that are identical or similar across a set of multiple structures. Without relying on topological constraints, `MHCXGraph` analysis can be focused on user-defined surface regions and support simultaneous analysis of multiple structures without requiring pairwise comparison. To optimize performance, the association graph is constructed using tokenized node triads embedded with discrete geometrical properties. This design enables comparison of input pMHC structures with different peptide lengths and MHC classes, including non-classical alleles, as well as cases involving non-canonical amino acids or even water molecules. Collectively, `MHCXGraph` provides a flexible and scalable framework for structure-based cross-reactivity analysis in immunological and therapeutic applications.

## 2 Results

### 2.1 MHCXGraph package

`MHCXGraph` is an open-source Python package that leverages graph-based operations to identify conserved exposed regions across multiple pMHC structures (Figure 1). It supports multiple execution modes and offers fully adjustable parameters, enabling flexible configuration to suit diverse user needs. For result analysis, the package provides an interactive dashboard that facilitates data exploration through graph visualizations and projections onto 3D structures.

**Figure 1.**
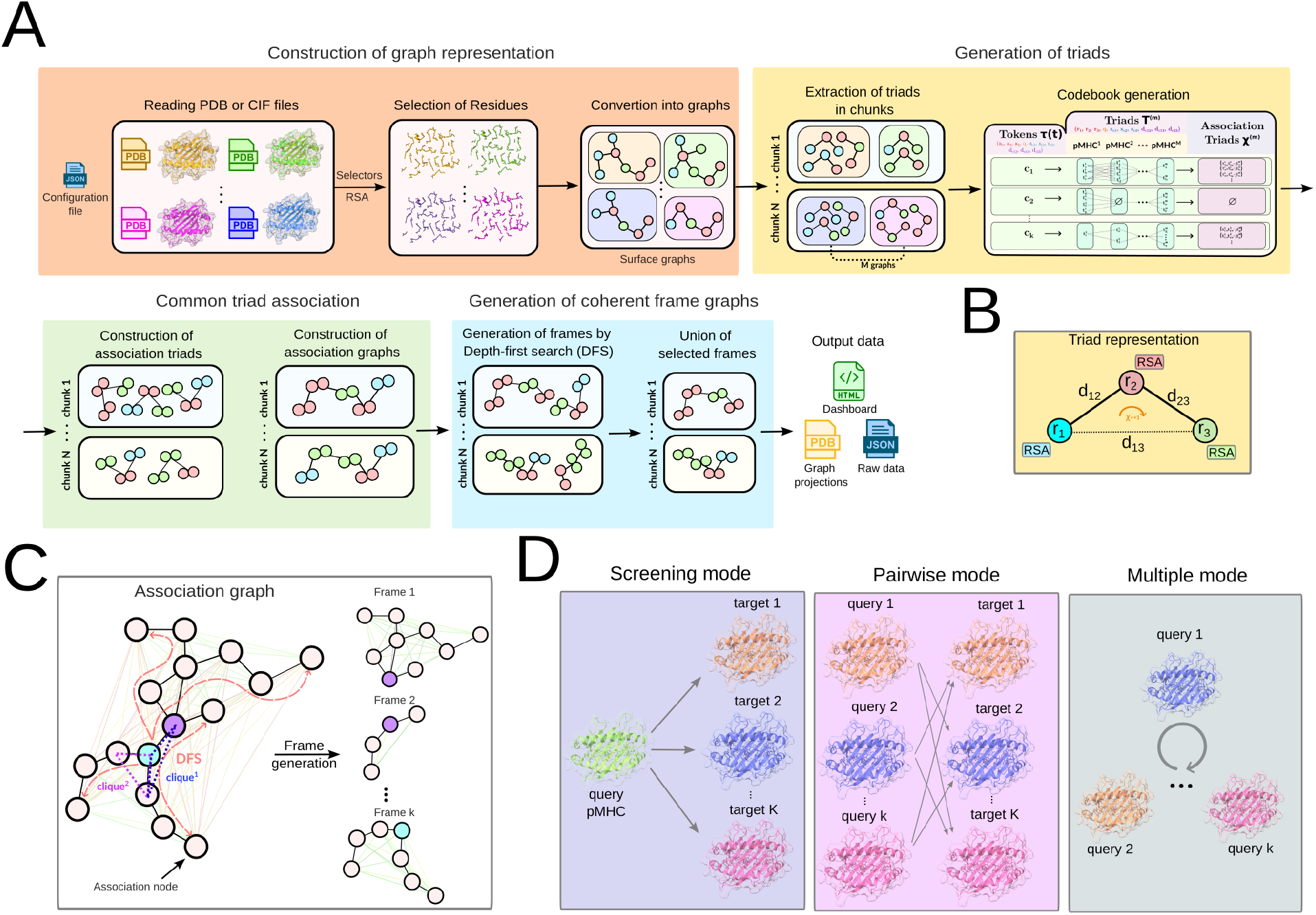
Schematic workflow of the `MHCXGraph` package. (A) Schematic workflow illustrating the main steps and algorithms of the method. (B) Detailed view of the triad representation, highlighting its features and attributes. (C) Detailed view of the coherent frame graph generation step, illustrating the operation of the depth-first search (DFS) algorithm on the association graph. Local clique searches supporting the DFS in identifying coherent nodes are also shown. Green and red lines connecting nodes indicate whether pairs of association nodes satisfy the coherence condition, i.e., whether all corresponding nodes respect the global distance threshold across input structures. (D) Schematic overview of the package execution modes.

#### 2.1.1 Algorithm and workflow

The package workflow consists of four main steps, detailed in the Methods section (Figure 1A). The first step is the construction of graph representations of pMHC surfaces named as a surface graph. In this step, selected solvent-exposed residues are represented as nodes, and edges are defined based on distance thresholds. To define appropriate thresholds for residue selection using relative solvent accessibility (RSA) and for edge construction, we performed a structural analysis of TCR:pMHC complexes, as detailed in the Supplementary Material (see section “Characterization of pMHC structural interfaces with TCRs”). For canonical residues, exposure is defined based on RSA threshold. Other molecules (e.g., ligands, water molecules, or non-canonical amino acids) can also be included in the graph representation, making it versatile. Instead of individual amino acids, users may choose to use amino acid classes based on physicochemical characteristics. Edges are constructed as undirected connections between nodes, defined by user-specified distance thresholds applied to the 3D coordinates of residues, according to the chosen node granularity. The selection of residues is an important step in the workflow, enabling users to focus the analysis on regions most relevant for TCR interaction. To support this process, we provide a curated list of pMHC residues that have already been observed to interact with TCRs in experimentally solved structures (58 positions for MHC-I and 35 for MHC-II; see section “Characterization of pMHC structural interfaces with TCRs” and Table 1 in the Supplementary Material). Additionally, users may opt to select residues based on secondary structure, such as the helical structures α1 and α2 of MHC, which contain important anchors for TCR interaction.

In the second step, each surface graph is decomposed into small, overlapping triads (three-node subgraphs), a critical step for computational efficiency. Each triad is encoded by a set of attributes, including inter-node distances, solvent accessibility, and chirality, capturing local structural features (Figure 1B). All these attributes are discretized into tokens, forming a codebook of triads that enables more efficient identification of associated triads across graphs. All steps starting from triad generation are performed in chunks to control memory usage.

The third step constitutes the core of the workflow and involves the association of common graph triads across the input graphs. Association requires matching triads from different input structures within the same token. This matching involves comparing node amino acid identities, triad chirality, inter-node distances, and differences in node RSA, with the latter two constrained by local distance and RSA difference thresholds defined by the users. This process results in the construction of an association graph containing residues shared among the input structures. Multiple association graph components, representing disconnected subgraphs, may be generated in this step.

However, even when local triad constraints are satisfied, association nodes from different triads can still present undesired distance deviations between them, indicating geometric incoherence. Therefore, in the fourth step, we extract fully coherent subgraphs, named frame graphs, from the association graph by selecting only association nodes in which all nodes satisfy a global distance threshold relative to other non-adjacent association nodes. This process is performed using a modified depth-first search (DFS) algorithm combined with local clique identification (Figure 1C). The result of this step is a set of frame graphs containing only fully coherent association nodes with consistent distances across all input structures.

#### 2.1.2 Running modes

`MHCXGraph` supports three running modes: Multiple, Pairwise, and Screening, all configurable via user-defined parameters 1D).

In multiple-analysis mode, all input structures are compared simultaneously, allowing the identification of conserved regions across the entire dataset. To ensure computational and memory efficiency, analyses involving multiple structures are performed in chunks. In pairwise-analysis mode, all non-redundant pairs of input structures are analyzed independently, which is useful for identifying conserved regions within subsets and for performing similarity comparisons between individual input structures. In screening mode, a reference structure is compared against a set of target structures. In addition, users can focus the analysis on the peptide and its immediate surrounding MHC environment.

#### 2.1.3 Visualization

`MHCXGraph` generates a diverse set of outputs to allow users to easily access and interpret the results. Standard outputs include HTML reports displaying graphs from different stages of the workflow, including surface graphs, association graphs, and frame graphs. Additionally, the package generates PDB files mapping the identified nodes onto the original 3D structures, as well as raw data files for downstream analysis.

By activating the dashboard visualization, users can access all outputs on a single interactive interface (Figure 2). The dashboard provides easy access to visualize graphs, map them onto 3D structures with full graphical control, and perform post-processing analyses.

**Figure 2.**
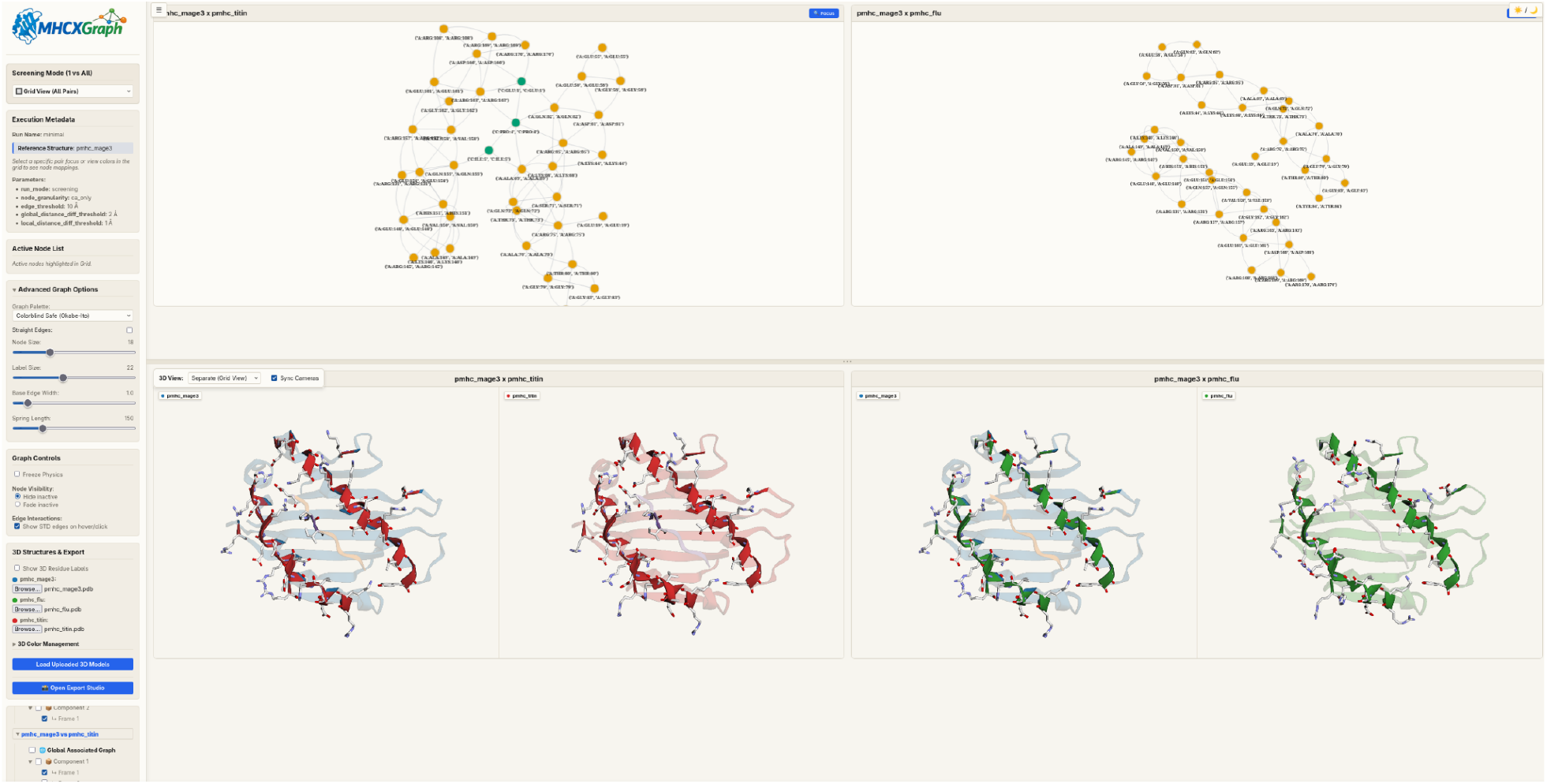
Illustrative example of the `MHCXGraph` dashboard. This example represents an output dashboard from an `MHCXGraph` run in screening mode. In this case, the MAGE-A3 melanoma-related peptide presented by HLA-A*01 is compared to a self-peptide derived from the titin protein and to a peptide derived from an influenza protein, both bound to the same HLA. The dashboard highlights, in the upper panel, the graph frames of the first component for each comparison, and in the bottom panel, the projection of the graphs onto the corresponding oriented 3D structures. On the left, a set of metadata and parameters can be accessed and visualized.

Importantly, although users may further process the raw data to define custom metrics, we provide a coverage-based similarity metric, a modified version of the metric described in [24], that quantifies the similarity between the graph representations of input proteins based on their coverage within the association graph.

### 2.2 Case studies

Here, we demonstrate the applicability and versatility of `MHCXGraph`, by evaluating the method across three representative case studies covering distinct biological contexts relevant to T cell recognition and cross-reactivity. These include: (1) the identification of conserved surface regions across prevalent HLA alleles, independent of bound peptides; (2) the analysis of TCR cross-reactivity among cancer-associated epitopes presented by the same MHC-I molecule; and (3) the investigation of cross-reactivity across different MHC-II alleles presenting HIV-derived peptides. Together, these case studies highlight the ability of MHCXGraph to detect structurally conserved and immunologically relevant features across diverse pMHC systems.

#### 2.2.1 Identifying conserved surface regions across multiple prevalent HLA alleles

The success of therapies targeting MHC molecules, such as TCR-based immunotherapies and emerging *de novo* MHC engager designs, may depend on the specific HLA allele expressed by an individual. As a first case study, we applied `MHCXGraph` to identify conserved residues on MHC surfaces across the most prevalent HLA alleles in human populations [25], independently of the bound peptide.

For this purpose, we selected 11 alleles—HLA-A01:01, HLA-A02:01, HLA-A03:01, HLA-A11:01, HLA-A24:02, HLA-B07:02, HLA-B08:01, HLA-B15:01, HLA-B40:02, HLA-B44:03, and HLA-C*07:02—representing three different loci (i.e., A, B, and C). All selected alleles had experimentally solved 3D structures available in the unbound state (i.e., not complexed with a TCR). Peptides from the structures were not considered in this analysis.

We executed `MHCXGraph` in pairwise mode, considering all non-redundant structure pairs. As selectors for graph construction, we used a predefined set of MHC-I residues known to interact with TCRs. A similarity heatmap was generated using a coverage-based index (see Methods section), defined as the fraction of unique nodes in association graph frames relative to the total number of nodes in both input graphs (Figure 3A). Hierarchical clustering showed that `MHCXGraph` grouped HLAs by gene locus, with a primary cluster containing HLA-A alleles and a second cluster comprising HLA-B and HLA-C alleles.

**Figure 3.**
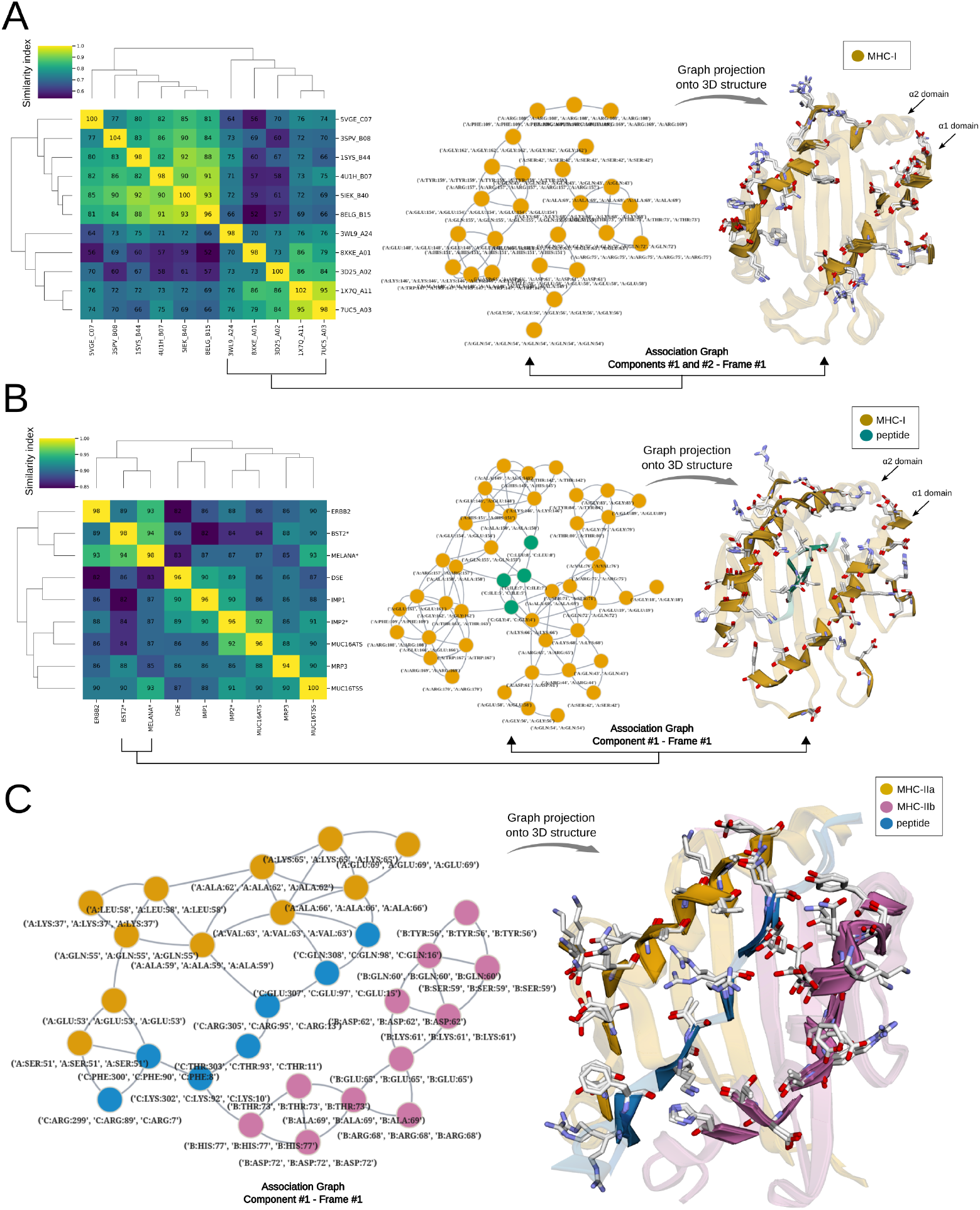
`MHCXGraph` analysis of different experimentally validated cross-reactivity case studies. (A) `MHCXGraph` analysis of 11 different HLAs without peptides, using a node granularity of Cα, an edge threshold of 10 Å, and local and global distance difference thresholds of 1.0 and 2.0, respectively. The heatmap on the left presents the pairwise comparison between each HLA based on the coverage-based similarity index. The values within each heatmap cell correspond to the number of unique nodes found across all frames of all components of the association graph. The diagonal represents twice the number of nodes in each input graph, corresponding to the maximum number of nodes that could be identified in a theoretical association graph of the same input protein. Dendrograms were built using the UPGMA hierarchical clustering method. In the center, the frames from the first two components of the association graph are shown. These originate from the multiple-analysis run mode applied to HLA-A only. The graph projection onto the superposed 3D structures is shown on the right, where nodes are depicted by residue side chains. (B) The same `MHCXGraph` analysis as in (A), but applied to a set of peptides presented by HLA-A*02. Asterisks on the heatmap labels (MELANA, BST2, and IMP2) indicate peptides that were determined to be recognized by the same Mel5 TCR. (C) `MHCXGraph` analysis in multiple-analysis run mode of cross-reactive HIV-1 peptides bound to different MHC-II alleles. Only the association graph and its projection onto the superposed 3D structures are shown. The node granularity used was the centroid of all residue atoms, with an edge threshold of 8.5 Å and with the use of the RSA feature in the triad. Local and global distance difference thresholds were set to 2. All analyses presented here were performed using an RSA threshold of 0.1 to define exposed nodes.

We then applied the multiple-analysis mode to the HLA-A subset to identify shared structural features. By analyzing the first graph frame from the first two association graph components (Figure 3A), we identified large common regions across the alleles, spanning the helices from both α1 and α2 domains. Projection of the graph onto the 3D structure provides a more detailed visualization of the conserved exposed residues. Despite this large conserved region, a non-conserved region can be observed at the end of the α1 helix, encompassing residues known to participate in TCR interactions in experimentally resolved complexes (Figure 3A; right panel). In the context of designing TCRs or protein binders, interactions with this region should be avoided when binding across all analyzed alleles is desired.

#### 2.2.2 Analyzing Mel5 TCR cross-reactivity in cancer-associated epitopes

As a second case study, we analyzed a set of 10-mer peptides presented by HLA-A*02, including three cancer-associated peptides shown to be recognized by the same Mel5 TCR [26]. In experimental T cell activation assays, peptides derived from the Melan-A (EAAGIGILTV), BST2 (LLLGIGILVL), and IMP2 (NLSALGIFST) proteins were identified as inducing a TCR response, with a weaker activation observed for the latter. Other peptides derived from ERBB2, DSE, IMP1, MRP3, MUC16ATS, and MUC16TSS did not show significant activation signals.

All peptides, whether cross-reactive or not, were modeled using the AlphaFold3 server [19], bound to HLA-A*02, and analyzed with `MHCXGraph` in pairwise mode. Clustering based on the similarity index identified the highest surface residue conservation between the two cross-reactive peptides derived from Melan-A and BST2, while IMP2 formed a more distant cluster, consistent with its reduced experimental activation.

The graph representation of the first frame from the first association graph component (Figure 3B) highlights conserved residues between the cross-reactive Melan-A and BST2 peptides. Structural mapping shows conservation in central peptide positions (e.g., GLY4, ILE5, and ILE7), which are known to participate in TCR interactions [26].

#### 2.2.3 Analyzing MHC-II Cross-Reactivity in HIV-Derived Epitopes

As a third case study, we investigated cross-reactivity across different MHC-II alleles (HLA-DR1, HLA-DR11 and HLA-DR15) presenting the most immunodominant HIV-1 capsid epitope (Gag293) and its shorter version RQ13. The complexes, HLA-DR1 and HLA-DR11 presenting the Gag293, and the HLA-DR15 presenting the RQ13, were experimentally shown to interact with the same TCR, named TCR F24 [27]. Experimental 3D structures of HLA-DR11–Gag293 and HLA-DR15–RQ13 (PDB IDs 6CPL and 6CPO, respectively) were obtained from the PDB, while the HLA-DR1-Gag293 was modeled with AlphaFold3 server [19].

Using multiple-analysis mode with RSA features in the triad representation and all atom node granularity, `MHCXGraph` identified seven conserved peptide residues, indicating a preserved binding register between the shorter RQ13 peptide and the full-length Gag293 peptide (Figure 3C). Additionally, conserved residues from both the α and β MHC-II domains were also identified, indicating an extensive interface for TCR cross-reactivity despite allele differences (Figure 3C).

#### 2.2.4 Runtime and memory usage benchmarking

To assess runtime and memory usage, we executed `MHCXGraph` in multiple and screening modes using two datasets, each comprising 100 pMHC structures. In the first dataset (Single HLA), all structures corresponded to the same MHC-I allele (HLA-A*02:01) bound to different peptides, whereas the second dataset (Diverse HLA) included different MHC-I alleles, each bound to distinct peptides. All experiments were performed on a workstation equipped with an Intel Core i7-14700 processor (2.10–5.40 GHz).

`MHCXGraph` analyzed 100 structures from both the Diverse and Single HLA datasets in a similar average runtime of 55 s, with peak memory usage of approximately 1.6 GB (Figure 4) in the multiple mode. A comparable performance profile was observed in screening mode, with an average runtime of 63 s for both datasets.

**Figure 4.**
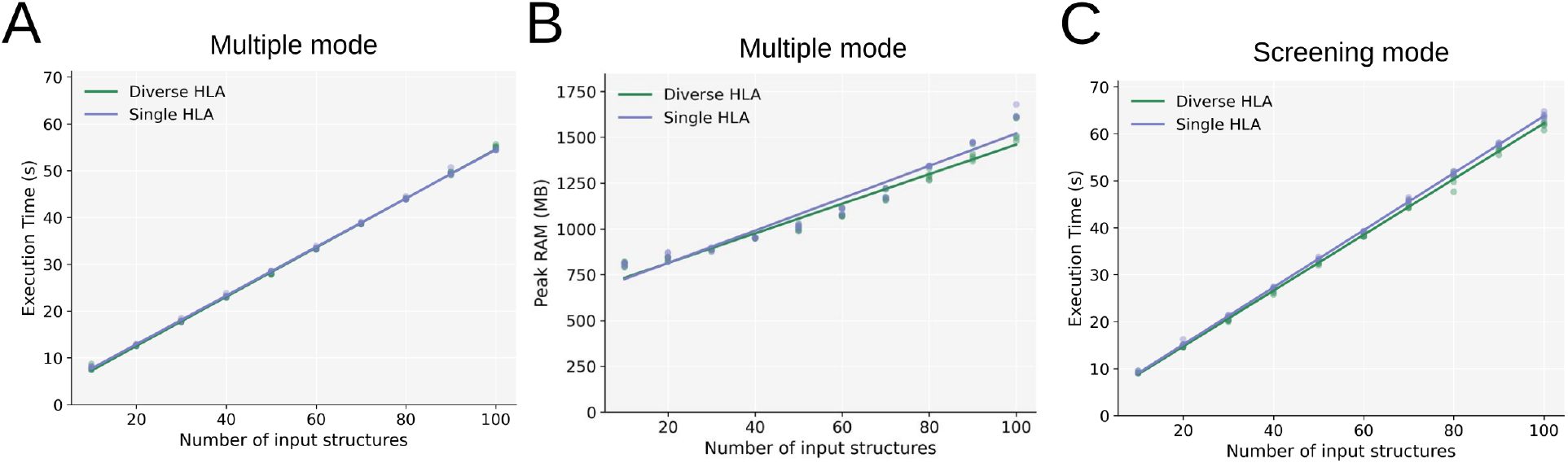
`MHCXGraph` runtime and memory usage analysis. (A) Runtime analysis of `MHCXGraph` in multiple mode using two datasets: the Diverse HLA dataset, comprising structures with different MHC-I alleles and peptides, and the Single HLA dataset, comprising structures from the same HLA-A*02:01 allele bound to different peptides. (B) Peak memory usage analysis of `MHCXGraph` in multiple mode. (C) Runtime analysis of `MHCXGraph` in screening mode. A linear trend line is presented in all analyses. For all analyses, `MHCXGraph` was executed using Cα node granularity and an edge threshold of 8.5 Å, without incorporating RSA into triad features. Local and global distance difference thresholds were set to 1 Å and 2 Å, respectively.

In both multiple and screening modes, runtime increased linearly as a function of the number of input structures within the evaluated range (*R*^2^ *>* 0.99). Based on the fitted linear model, runtime increased by approximately 0.5 s and 0.6 s per additional pMHC structure in multiple and screening modes, respectively. In contrast, peak memory usage followed a less strictly linear trend compared to runtime, particularly for larger numbers of input structures in multiple mode (*R*^2^ = 0.95 on average). On average, memory usage increased by approximately 8.5 MB per additional pMHC structure.

## 3 Discussion

The natural recognition of multiple MHC-presented peptides by a single TCR, despite being an essential process for the recognition of a broad array of antigenic peptides by the TCR repertoire, is one of the main challenges when developing TCR-guided therapies [28, 8, 7]. The understanding of similarities between peptides presented by MHC is also relevant in the context of vaccines based on T-cell responses [11]. For instance, in the development of pan-viral vaccines, one may want to select similar antigenic peptides for MHC presentation while ensuring their divergence from self-peptides [29].

The structural variability of peptide-exposed residues involved in TCR engagement, including register shifting [30], and the sequence and functional variability of MHC alleles presenting these peptides to TCR-specific populations [31], which may or may not lead to TCR cross-reactivity, create a particular need for structure-based comparison methods that sequence-based approaches alone cannot fully address [20].

In this context, we presented `MHCXGraph`, a graph-based method that can identify common regions across pMHC surfaces. `MHCXGraph` has several advantages and functionalities: it is alignment-free, it can focus on the most relevant TCR-interacting regions, it can be applied not only to the most common MHC-I but also to other MHC-like structures with or without peptide, and it can be directly used to analyze pairs or multiple structures.

The basis of the `MHCXGraph` algorithm is the maximum common subgraph problem for detecting subgraph isomorphism, which has previously been addressed to identify surface or structural similarities between proteins [24, 22, 21]. In classical approaches, an association graph is first built from the product graph of two input graphs, followed by a maximum clique operation to detect the common subgraph [24]. Here, we employ a related strategy, but with key differences to improve computational performance and geometric accuracy. These include, for example, performing the Cartesian product at the level of tokenized triads and generating coherent subgraphs through a DFS algorithm coupled with a local clique search. The latter aims to avoid the computationally expensive (NP-hard in general) maximal clique problem. Comparing to individual node associations, the use of triads as the basic unit of association introduces stronger geometric constraints, capturing local structural context beyond individual residues and reducing spurious matches. Despite its advantages, the use of triads restricts the detection of common surface regions to those involving at least three connected residues. However, this is not a limitation for MHC interaction interfaces, which are typically large and comprise, on average, 18 residues, as shown in our structural analysis.

Although MHCXGraph has been applied to analyze hundreds of structures, as demonstrated in the computational time analysis, and can scale linearly to thousands of structures within minutes in screening mode, its structure-based nature makes it particularly suitable as a final step in cross-reactivity detection workflows. Starting from a set of pMHC candidates derived, for example, from large-scale sequence-based proteome analyses, such as those implemented in the `ARDitox` pipeline [32], MHCXGraph can identify conserved surface regions and provide highly accurate structural validation and interpretability for cross-reactivity analysis.

Through different case studies, we demonstrated the usability of the method. `MHCXGraph` was able to identify conserved surfaces shared by pMHCs recognized by the Mel5 TCR [26] and cluster them in comparison with non-cross-reactive cases. It was also capable of identifying shared regions across different HLA alleles, even without the bound peptide, and grouping them according to their similarity at the TCR interface. While the identification of conserved pMHC regions can be useful for avoiding cross-reactivity, the identification of conserved sites across multiple alleles can be useful when designing a pan-allelic binder and regulating alloselectivity [33]. Finally, we also demonstrate the capability of the method to work with MHCs other than classical MHC-I, such as MHC-II, which, through an open groove at both ends, presents peptides with a broader range of lengths and flexible flanking segments [34].

Although `MHCXGraph` was developed with a focus on pMHCs and involved extensive testing in this context, the graph-based approach presented here can be directly applied to any type of protein surface. A practical application not addressed in this study is the identification of conserved regions on the surfaces of different TCR structures. When focused on TCR CDRs, the method could identify shared residues across clonally expanded T-cell populations with TCRs that recognize the same pMHC [35].

Results from large `MHCXGraph` analyses may be useful for addressing currently challenging TCR- and pMHC-related predictive tasks. The structural clustering of pMHC surfaces with `MHCXGraph` could improve the separation between training and test sets in machine learning models aimed at predicting TCR interactions with unseen peptides [36]. The identification of poorly conserved pMHC surfaces by `MHCXGraph` could also assist in assigning better TCR:pMHC negative pairs generated from positive shuffling [37, 38]. Ultimately, high-quality data generated by `MHCXGraph` could be used to train novel machine learning models for predicting protein surface similarities.

In conclusion, `MHCXGraph` package provides a versatile approach for studying, detecting, and interpreting potential cross-reactivity of peptides presented by MHC using structural information encoded in graph representations. MHCXGraph is expected to be integrated into computational pipelines for cross-reactivity discovery and to contribute to the development of safer TCR-based therapies and vaccines.

## 4 Methods

### 4.1 Package implementation and execution

The `MHCXGraph` package is executed using a JSON configuration file (`manifest.json`) that defines a set of entries controlling its behavior. Key parameters include input structures (`inputs`), residue selectors (`selectors`), edge threshold `(edge_threshold)`, node granularity `(node_granularity)`, and Relative Solvent Accessibility (RSA) filtering `(rsa_filter)`. A comprehensive list of parameters and a sample manifest file are available in the Supplementary Material (Table 2 and “Example of Manifest file” section, respectively). Additionally, advanced parameters are described in the following sections.

Structure inputs can be provided in either PDB or CIF file formats, either as a directory or as individual file paths. The minimal structural preprocessing recommended is the removal of TCRs or other macromolecule binders, as these can affect residue accessibility. Residue or chain selection can be specified using the `selectors` parameter. A predefined list of potential TCR-interacting residues, derived from structural analysis of TCR:pMHC complexes, is available (Supplementary Material, Table 1); however, its use requires MHC structures to follow IMGT numbering, which can be achieved using a preprocessing subcommand included in `MHCXGraph` package. Alternatively, residues may be selected based on secondary structure (e.g., helices), which is particularly useful when input structures do not follow a conventional numbering scheme but the user still wishes to focus on MHC regions most relevant to TCR interactions, such as the α1 and α2 helices. Optional parameters (`include_waters`, `include_ligands`, and `include_noncanonical_residues`) allow inclusion of non-protein entities (e.g., water, ligands, and modified amino acids).

All processes related to reading, processing and writing of 3D structures in `MHCXGraph` are handled using the Biopython package [39], while the secondary structure assignment is performed using the `PyDSSP` implementation of `DSSP` [40]. Instead of working with individual amino acids, users may choose to group them into classes based on physicochemical descriptors, enabling the identification of similar, rather than strictly identical, regions.

#### 4.1.1 Surface Graph Generation

Each pMHC structure is converted into a graph representation, referred to as surface graph. Let *M* be the number of input graphs (i.e., input pMHC structures), then *G*^(*m*)^ = (*V* ^(*m*)^, *E*^(*m*)^), ∀*m* ∈ {1, …, *M*}, where nodes *V* ^(*m*)^ denotes the set of nodes, being node *v* ∈ *V* ^(*m*)^ defined as *v* = (*c, a, n*), where *c* denotes the chain identifier, *a* the amino acid name (or amino acid class), and *n* the residue number. Edges *E*^(*m*)^ represent spatial proximity relationships defined based on a Euclidean distance threshold (`edge_threshold`).

Node coordinates may be computed using different atom representations (e.g., all-atom centroid, side-chain centroid, backbone centroid, or Cα atom). Solvent exposure is determined for each node corresponding to canonical amino acids using Relative Solvent Accessibility (RSA), computed via the Lee–Richards algorithm as implemented in the FreeSASA Python package [41], and normalized using the Wilke scale [42]. For non-canonical residues, water molecules, and ligands, node coordinates consider all atoms and the accessible surface area (ASA) is used, providing an additional structural descriptor for these nodes.

From each surface graph, an induced subgraph *G*^*′*(*m*)^ is then constructed based on user-defined residue selection of residues. Given a subset of nodes *V* ^*′*(*m*)^ ⊆ *V* ^(*m*)^, defined according to residue identity, chain, or secondary structure, and filtered by solvent exposure, the resulting subgraph is the induced graph *G*^*′*(*m*)^ = (*V* ^*′*(*m*)^, *E*^*′*(*m*)^), where *E*^*′*(*m*)^ = {(*u, z*) ∈ *E*^(*m*)^ | *u, z* ∈ *V* ^*′*(*m*)^}.

Graph construction and attribute assignment are performed using NetworkX [43] package. Default thresholds for RSA and edge definition were derived from structural analyses of TCR:pMHC interfaces (see Supplementary Material; “Characterization of pMHC structural interfaces with TCRs” section).

#### 4.1.2 Triad Graph Generation

Each surface graph is then decomposed into sets of overlapping node triplets (i.e., triad graphs). A triad is defined as an ordered triplet of nodes *t* = (*v*_*i*_, *v*_*j*_, *v*_*k*_), where nodes *v*_*i*_, *v*_*j*_, *v*_*k*_ ∈ *V* ^*′*(*m*)^ and are connected in the underlying graph structure. Each graph *G*^*′*(*m*)^ generates a set *T* ^(*m*)^ of triads. To ensure consistency, the outer nodes of each triad are sorted alphabetically. When the outer nodes share the same amino acid identity, they are further ordered based on their distances to the central node, ensuring a unique and reproducible representation.

Graph triads *t* ∈ *T* ^(*m*)^ are encoded using a feature mapping *ϕ* : *T* ^(*m*)^ → ℱ, defined as *ϕ*(*t*) = (*r*_1_, *r*_2_, *r*_3_, *q, s*_1_, *s*_2_, *s*_3_, *d*_12_, *d*_13_, *d*_23_), where *r*_*i*_ are the residue identifiers (chain:resname:resnumber), *q* encodes chirality, *s*_*i*_ are RSA values, and *d*_*ij*_ are pairwise Euclidean distances between nodes *i* and *j* in the triad.

Triad chirality *q* is encoded as a discrete value (+1 or −1) computed using backbone and side-chain geometric information, as described in the Supplementary Material (see “Determination of chirality of node triads” section). In this procedure, glycine residues are assigned a virtual Cβ atom, and triads involving other than canonical amino acids are assigned a chirality value of 0. This descriptor is invariant to rigid transformations but sensitive to mirror reflections, enabling discrimination between enantiomeric configurations of triads.

To focus on specific chains, such as peptides in pMHC complexes, users can activate the (`filter_triads_by_chain`) option to retain only triads containing at least one node from the selected chain. Unlike restricting the analysis to a single chain, this approach preserves neighboring nodes from other chains, allowing the identification of structural similarities in the local environment of the target chain.

For efficient comparison, triads *t* are discretized into a finite token space, defining a codebook of triad classes. This is achieved by mapping continuous features, such as inter-residue distances and solvent accessibility, into discrete bins via a function *τ* : *T* ^(*m*)^ → 𝒞, where each triad is associated with a token *τ* (*t*) = (*a*_1_, *a*_2_, *a*_3_, *q, s*_*c*1_, *s*_*c*2_, *s*_*c*3_, *d*_*c*12_, *d*_*c*13_, *d*_*c*23_), in which *a*_*i*_, denote amino acid identities or classes, and *s*_*ci*_, *d*_*cij*_ represent discretized RSA and distance values, respectively.

The number of bins is defined by the user through the choice of bin width (`distance_bin_width`) and the edge threshold (`edge_threshold`). Smaller bin widths increase the number of classes and enforce stricter similarity criteria. Although discretization enables efficient comparison, it introduces hard bin boundaries that may affect values near bin edges. To mitigate this hard-bin effect, a given distance or RSA value can be assigned to multiple bins based on the `local_distance_diff_threshold` parameter, which defines the maximum tolerated distance when comparing corresponding node distances (*d*_12_, *d*_13_, or *d*_23_) between triads.

After discretization, the set of triads *T* ^(*m*)^ is partitioned according to their token assignments. In particular, the mapping *τ* : *T* ^(*m*)^ → 𝒞 induces a decomposition of *T* ^(*m*)^ into subsets of structurally comparable triads. Formally, for each token *c* ∈ 𝒞, we define the subset of triads in graph *G*^*′*(*m*)^ associated with this token as 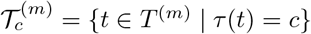.

Each set 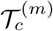 thus groups triads that share the same discretized geometric and physicochemical descriptors. This organization restricts subsequent comparisons to triads that are already compatible at the token level, avoiding unnecessary cross-comparisons between structurally dissimilar elements.

As a result, the matching process can be carried out independently within each token class *c*, which substantially reduces the combinatorial complexity of the association step and prepares the construction of association triads via restricted Cartesian products across graphs.

#### 4.1.3 Association triad generation and association graph construction

Triads from different input graphs are combined into association triads using a restricted Cartesian product within each token. For a given token *c* ∈ 𝒞, we consider the sets 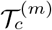 previously defined for each graph *G*^*′*(*m*)^ and restrict attention to tokens that are represented in all input graphs, i.e., such that 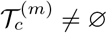 for all *m* ∈ {1, …, *M*}. For such tokens, the set of association triads, is given by 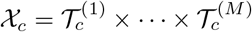 and an association triad is defined as an element of 𝒞_*c*_ as follows *t*′ := (*t*^(1)^, …, *t*^(*M*)^), with 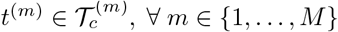, representing one triad selected from each input graph, all sharing the same token assignment. This construction ensures that only geometrically and physicochemically comparable triads are combined. In practice, the Cartesian product 𝒳_*c*_ is not computed exhaustively. Instead, candidate tuples are generated incrementally and filtered during construction using bounds on continuous descriptors, reducing combinatorial complexity.

To eliminate potential false positives introduced during discretization and multiple class assignment, association triads are further filtered by enforcing consistency in their continuous descriptors. This filtering is defined by a function Ψ : *𝒳*_*c*_ → {0, 1}, such that Ψ(*t*^(1)^, …, *t*^(*M*)^) = 1 ⇔ ∀*m* ∈ {1, …, *M*}, ∀(*i, j*) ∈ {(1, 2), (1, 3), (2, 3)}, 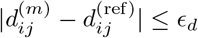 and 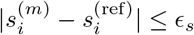, where ref denotes a reference input (selected as the graph with the fewest triads), and *ϵ*_*d*_ and *ϵ*_*s*_ are user-defined thresholds for distances and RSA values, respectively. The set of accepted association triads for a given token *c* ∈ 𝒞 is defined by: 𝒜_*c*_ = {*t*′ ∈ 𝒞_*c*_ | Ψ(*t*′) = 1}.

To reconstruct a unified graph structure from association triads, node correspondences are extracted from the accepted association triads, and edges are defined between nodes that co-occur within the same accepted association triad. Formally, for each accepted association triad *t*′ = (*t*^(1)^, …, *t*^(*M*)^), *t*′ ∈ 𝒜_*c*_, where for each 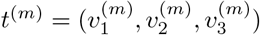, we create three association nodes 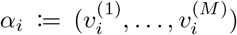, for *i* ∈ {1, 2, 3}. The set of association nodes *V*_*A*_ is then given by all such tuples *α*_*i*_ induced by accepted association triads. Edges are added between association nodes corresponding to adjacent nodes in the ordered triads, namely (*α*_1_, *α*_2_) and (*α*_2_, *α*_3_) for each *t*′ ∈ 𝒜_*c*_. The resulting association graph is defined as *A* = (*V*_*A*_, *E*_*A*_). This process may yield multiple disconnected graph components, each corresponding to a distinct association graph component.

#### 4.1.4 Generation of coherent frame graphs

The creation of association graphs, as described above, introduces an important limitation: a potential loss of coherence between non-adjacent distant association nodes. Here, coherence refers to the property by which a corresponding pair of residues, represented as nodes from multiple structures, exhibits similar Euclidean distances across all inputs (Figure 1). Formally, for two association nodes *α*_*i*_, *α*_*j*_ ∈ *V*_*A*_, coherence requires that the pairwise distances between their corresponding residues are consistent across all input graphs, i.e., 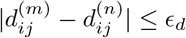 for all *m, n* ∈ {1, …, *M*}.

During association graph construction, this property is guaranteed only locally, as coherence is verified solely for residue pairs involved in matched triads and for edges introduced during triad expansion. Consequently, while adjacent association nodes satisfy local geometric consistency, no such guarantee is imposed on distant, non-adjacent nodes within the same connected component.

To enforce global consistency, coherence is evaluated for all unordered pairs of association nodes. Let *N* = |*V*_*A*_| be the number of association nodes. Since pairwise distance matrices are symmetric, i.e., 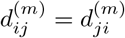, it is sufficient to consider only unordered pairs (*i, j*) with *i < j*. The total number of such unique pairs is therefore 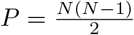. For each node pair (*i, j*), we define the vector of distances across the *m* input structures as 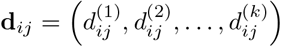, where 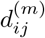 denotes the Euclidean distance between the residues corresponding to *α*_*i*_ and *α*_*j*_ in structure *m*. Rather than storing *M* full distance matrices of size *N × N*, we exploit symmetry and represent the data in compact form as a matrix *D* ∈ ℝ^*M ×P*^, where each column corresponds to a unique unordered pair (*i, j*). This representation preserves all pairwise geometric information while avoiding redundant storage.

Global coherence is then defined by bounding the variation of distances across structures. A pair (*i, j*) is considered coherent if max **d**_**ij**_ −min **d**_**ij**_ ≤ *ϵ*_*g*_, where *ϵ*_*g*_ is a user-defined threshold given by `global_distance_diff_threshold`. In practice, this threshold may be specified globally or made dependent on the chain identity of the residues involved, allowing different tolerance values to be applied to different combinations of chains. This induces a Boolean coherence relation *C*(*i, j*) defined as *C*(*i, j*) = 1 if max **d**_**ij**_ − min **d**_**ij**_ ≤ *ϵ*_*g*_, and *C*(*i, j*) = 0 otherwise.

Given an association graph component *A* = (*V*_*A*_, *E*_*A*_) and the coherence relation *C*(*i, j*), coherent frame graphs are constructed as subgraphs that satisfy both adjacency and global geometric consistency constraints. Thus, a frame graph is an induced subgraph *F* = (*V*_*F*_, *E*_*F*_), with *V*_*F*_ ⊆ *V*_*A*_, such that *E*_*F*_ = {(*α*_*i*_, *α*_*j*_) ∈ *E*_*A*_ | *α*_*i*_, *α*_*j*_ ∈ *V*_*F*_} and *C*(*i, j*) = 1, ∀*i, j* ∈ {1, …, *N*} with *α*_*i*_, *α*_*j*_ ∈ *V*_*F*_, *i* ≠ *j*. That is, all node pairs in the frame are mutually coherent across all input structures.

To identify such subsets, we define the coherence graph *G*_*C*_ = (*V*_*A*_, *E*_*C*_), *E*_*C*_ = {(*α*_*i*_, *α*_*j*_) | *C*(*i, j*) ≠ 1, *i j*} in which edges represent pairwise coherence. A subset of nodes is mutually coherent if and only if it forms a clique in *G*_*C*_.

Frame generation is performed independently for each connected component of the association graph. Nodes are first ordered according to their degree, and each node is used as an initial seed. For a given seed node, its adjacency neighborhood in *A* defines a candidate set. Since adjacency alone does not guarantee mutual coherence, a coherence graph is implicitly constructed over this candidate set, and maximal cliques are extracted using the Bron-Kerbosch algorithm [44]. Each clique represents a subset of nodes that are pairwise coherent and serves as a valid seed for frame construction.

Once coherent seed nodes are identified, frame expansion is performed using array-level operations derived from the coherence matrix. At each step, candidate nodes are restricted to those that are both adjacent in *A* and coherent with all nodes currently in the frame. This restriction is efficiently implemented by intersecting the corresponding rows of the coherence relation, avoiding explicit pairwise comparisons.

The traversal of candidate nodes follows a depth-first search (DFS) strategy (Figure 1), iteratively expanding each frame while preserving both adjacency and global coherence constraints. Expansion continues until no additional nodes satisfy these conditions.

When expansion terminates, the resulting subgraph is evaluated as a candidate frame. Valid frames must satisfy structural constraints, including a minimum number of edges and the absence of isolated nodes. Finally, redundant frame graphs are removed by eliminating subgraphs that are strict subsets of larger frames. The remaining maximal subgraphs define the final set of coherent frame graphs.

#### 4.1.5 Processing in chunks

Despite the efficient implementation of the method and extensive use of array operations, increasing the number of protein inputs processed simultaneously can introduce computational challenges in terms of memory usage and execution time on conventional systems, particularly during association triad construction and frame graph generation. To address these limitations, the algorithm processes input proteins in chunks, as specified by the user via the `max_chunks` parameter. Larger chunk sizes enable more proteins to be processed simultaneously but require increased memory resources. Chunks are currently processed sequentially: the first group of protein inputs is processed, followed by subsequent groups. Each chunk yields a set of frame graphs. If, after the initial processing layer, the number of resulting graphs still exceeds the maximum number of inputs allowed per chunk, an additional layer of chunking is applied. This process is repeated iteratively until completion.

Importantly, a given component within a chunk may generate multiple frame graphs containing redundant association nodes. To prevent the propagation of redundant information to subsequent chunk-processing layers, a union operation is applied to merge all frames into a single graph based on node identifiers, using NetworkX. Although this step may temporarily reintroduce incoherent nodes, this issue is resolved in the final chunk-processing layer, where the union operation is no longer applied and coherence is fully enforced.

#### 4.1.6 MHCXGraph output

`MHCXGraph` can generate a diverse set of outputs to help users visualize, interpret, and analyze the results. Output types can be selected based on user preferences. One type of output consists of HTML files containing interactive graph representations generated during execution. These visualizations include the input graphs, the association graphs, the association component graph, and the frame graphs generated for each component. Another output option consists of PDB files in which the input structures are superposed based on the residues corresponding to the association and frame graphs, facilitating visual inspection of the results in molecular visualization software. In addition, individual PDB files of the input structures are generated with the association and frame graph information embedded. `MHCXGraph` also outputs CSV files containing the raw data of the association nodes found, which can be used for post-processing and data analysis.

Although users have full control over output post-processing and analysis metrics, we define and provide a coverage-based similarity index derived from the coherent frame graphs. This metric represents the fraction of nodes from both graphs that are recovered in at least one coherent matched region. Let *G*^(*m*)^ = (*V* ^(*m*)^, *E*^(*m*)^) and *G*^(*n*)^ = (*V* ^(*n*)^, *E*^(*n*)^) be two input graphs, and let ℱ denote the set of all frame graphs obtained after the frame generation step. We define *U* ^(*m*)^ ⊆ *V* ^(*m*)^ and *U* ^(*n*)^ ⊆ *V* ^(*n*)^ as the sets of nodes from *G*^(*m*)^ and *G*^(*n*)^, respectively, that appear in at least one frame ℱ graph in . The coverage-based similarity index is then computed as 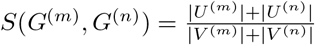.

Users can also opt to generate a dashboard where they can assess all `MHCXGraph` outputs on a single interactive visualization page that can be accessed offline (Figure 2). The dashboard can also be used to analyze results and generate high-quality images.

#### 4.1.7 Renumbering of the amino acid sequence in MHC structures

In order to enable users to identify TCR-contacting residues on the MHC and thus restrict the graph search space to only MHC residues observed to interact with TCRs, we provide an accessory program that renumbers the input MHC structure according to the IMGT domain numbering scheme [45]. This program must be executed prior to graph analysis if the user wants to guarantee full compatibility between the list of provided MHC hotspot residues and the MHC structure numbering (Supplementary Material, Table 1).

The renumbering algorithm relies on sequence alignment between the query amino acid sequence of the MHC structure and a list of gapped MHC template sequences (https://www.imgt.org/IMGTrepertoireMH/Proteins/protein/G-DOMAIN/Gdomains.html). The numbering follows the IMGT unique numbering scheme for MHC groove G-DOMAIN and G-LIKE-DOMAIN (https://www.imgt.org/IMGTScientificChart/Numbering/IMGTGsuperfamily.html). MHC-I is treated as two separate domains, α1 and α2, while MHC-II consists of two separate chains, α and β.

The program uses Python scripts built with Biopython and first identifies and extracts MHC-I or MHC-II sequences from the input structures. The extracted sequences are then aligned against all available templates in the ungapped format using global pairwise sequence alignment. A ranking is defined primarily based on the alignment score. In the case of structural gaps in the input structures, residue gaps are inserted into the query sequence representation to account for missing positions, provided that a numbering gap exists in the structure. The pairwise sequence alignment is then projected onto the IMGT numbering positions based on the selected gapped template. Insertions relative to the template are assigned to numbered gap columns when available. In the algorithm, only the MHC sequence in the structure is affected by the renumbering.

#### 4.1.8 3D modeling of pMHC structures

For case studies of `MHCXGraph` applications, pMHC sequences were modeled using the AlphaFold3 server [19] when experimental structures were not available. Only the top-ranked AlphaFold model was used for analysis.

To evaluate computational time and memory usage of `MHCXGraph`, we generated two separate datasets of modeled pMHC structures, each containing 100 entries. The first dataset comprised 100 distinct peptides bound to the same HLA-A*02:01 allele, whereas the second included 100 distinct peptides bound to different MHC-I alleles. All pMHC sequences were obtained from the VDJdb database [46], selecting entries with a confidence score of at least 2 and peptide lengths of 9 or 10 residues. Structures were modeled using ColabFold (v1.5.5) [47], and only the top-ranked models were retained for analysis. Only structures in which the peptides were appropriately bound within the MHC groove in a canonical register were considered.

## 5 Funding

This study was supported by the Serrapilheira Institute (grant number Serra–R-2401-47149). This study was financed, in part, by the São Paulo Research Foundation (FAPESP), Brasil, according to the following grants: #2024/12890-5, São Paulo Research Foundation (FAPESP) (to H.V.R.F); #2025/19395-2, São Paulo Research Foundation (FAPESP) (to R.L.B.R.M.); #2025/00373-9, São Paulo Research Foundation (FAPESP) (to C.D.M.S.S); #2024/20196-1, São Paulo Research Foundation (FAPESP) (to S.C.A).

The opinions, hypotheses, and conclusions or recommendations expressed in this material are the responsibility of the author and do not necessarily reflect the views of FAPESP. The funders had no role in study design, data collection and analysis, decision to publish, or preparation of the manuscript.

## 6 Acknowledgements

We are grateful to the Brazilian Biosciences National Laboratory (LNBio), part of the Brazilian Center for Research in Energy and Materials (CNPEM), a private non-profit organization under the supervision of the Brazilian Ministry of Science, Technology, and Innovations (MCTI), for providing staff and infrastructure for the development of this study. This research used the facilities and the Marvin HPC cluster of the Brazilian Biosciences National Laboratory (LNBio). The staff of EDB is acknowledged for their assistance during the experiments (#20242773). We thank José Geraldo de Carvalho Pereira (EDB/CNPEM) for valuable discussions on algorithm development. We also especially thank Dr. Brian Pierce (University of Maryland) for insightful methodological and immunological discussions. Finally, we thank the students of the ILUM School of Science for their contributions to the early development of preliminary algorithms used in this work.

## 7 Data Availability

The `MHCXGraph` source code is publicly available at https://github.com/cnpem/MHCXGraph under AGPL 3.0 license and can be installed via PyPI (https://pypi.org/project/MHCXGraph). The documentation and tutorial are available at https://cnpem.github.io/MHCXGraph/.

## Notes

https://github.com/cnpem/MHCXGraph/

### Competing Interest Statement

The authors have declared no competing interest.

