## Supplementary Material for "MHCXGraph: A Graph-Based approach to detecting T cell receptor cross-reactivity"

A PREPRINT

**Carlos Daniel Marques Santos Simões** [1,2], **Rocio Lucia Beatriz Riveros Maidana** [1], **Samuel Chagas de Assis** [1,2], **Joao Victor da Silva Guerra** [1], and **Helder Veras Ribeiro-Filho** [1,2,*]

[1]Brazilian Biosciences National Laboratory, Brazilian Center for Research in Energy and Materials, Campinas, São Paulo, Brazil

[2]Graduate Program in Pharmaceutical Sciences, Faculty of Pharmaceutical Sciences, University of Campinas, Campinas, São Paulo, Brazil

April 7, 2026

#### 1 Characterization of pMHC structural interfaces with TCRs

##### 1.1 Construction of the TCR:pMHC structure dataset for structure analysis

A set of TCR structures bound to pMHC was collected from the TCR3d database [1] on June 23, 2025, without restriction to species. A total of 292 MHC-I and 96 MHC-II PDB entries were retrieved, and only structures with resolution better than 2.5Å were retained, resulting in 97 MHC-I and 15 MHC-II structures selected for further analysis. PDB ID 9EJI (an MHC-II structure) was excluded due to its unconventional binding mode, which lacks contacts with the peptide. PDB ID 2VLJ was also excluded as it is redundant with PDB ID 1OGA, which has a higher resolution.

To ensure compatibility in residue numbering across MHC structures, each collected structure was renumbered according to a IMGT numbering scheme as described in this Supplementary Material.

The bound complexes were analyzed to identify pMHC residues in contact with the TCR, using a 5Å distance threshold between all atoms. For each PDB, contact residues were determined and then aggregated to generate a list of MHC-I and MHC-II residues observed to interact with the TCR in at least one analyzed structure. In structure analysis, the RSA was computed with the DSSP program (version 4.0.4), using the Wilke scale [2].

##### 1.2 Characterizing pMHC structural interfaces with TCRs to support MHCXGraph analysis

To support MHCXGraph analysis and parameter selection, we first represented and characterized, from a graph perspective, the pMHC (Class I and Class II) interfaces interacting with TCRs in solved 3D complex structures.

From the analysis of 97 MHC class I and 15 MHC class II structures in complex with TCRs from the PDB, we identified 58 distinct MHC positions for MHC-I and 35 for MHC-II that make contact with the TCR in at least one complex, considering a 5Å distance cutoff . These positions are provided to the user as input residue selectors, representing target graph nodes when constructing surface graphs in the MHCXGraph pipeline. This allows the method to focus on the most important MHC regions involved in interactions with the TCR. Importantly, only a subset of these positions constitutes the interaction interface in any given pMHC complex: on average, 18 residues for MHC-I and MHC-II. Peptide residues contribute at least 4 in pMHC-I and 6 in pMHC-II.

| MHC | Positions (IMGT numbering) |
|---|---|
| MHC-I | 18, 19, 42, 43, 44, 54, 55, 56, 58, 59, 61, 62, 63, 64, 65, 66, 68, 69, 70, 71, 72, 73, 75, 76, 77, 79, 80, 83, 84, 89, 108, 109, 142, 143, 145, 146, 147, 148, 149, 150, 151, 152, 153, 154, 155, 156, 157, 158, 159, 161, 162, 163, 165, 166, 167, 169, 170, 171 |
| MHC-IIα | 37, 51, 52, 53, 55, 56, 58, 59, 60, 62, 63, 65, 66, 67, 69 |
| MHC-IIβ | 56, 57, 59, 60, 61, 62, 63, 65, 66, 67, 68, 69, 70, 71, 72, 73, 74, 77, 78, 81 |

Table 1: List of MHC-I and MHC-II positions at the TCR interface. This list includes positions from MHC-I and MHC-II (alpha and beta chains) that were observed in at least one experimentally solved TCR:pMHC complex to be in contact with the TCR (within 5Å distance). Importantly, all analyzed structures were renumbered according to the IMGT numbering scheme.

The average Relative Solvent Accessibility (RSA) of pMHC residues that are in contact with the TCR (5Å distance), when analyzing the corresponding unbound pMHC structures (i.e., without the TCR), was 0.3 for both MHC-I and MHC-II cases (Supplementary Figure S1A and S1B). The RSA threshold of 0.1 lies at approximately the 85th percentile for both MHC classes, indicating that it is a suitable threshold for selecting nodes to MHCXGraph analysis.

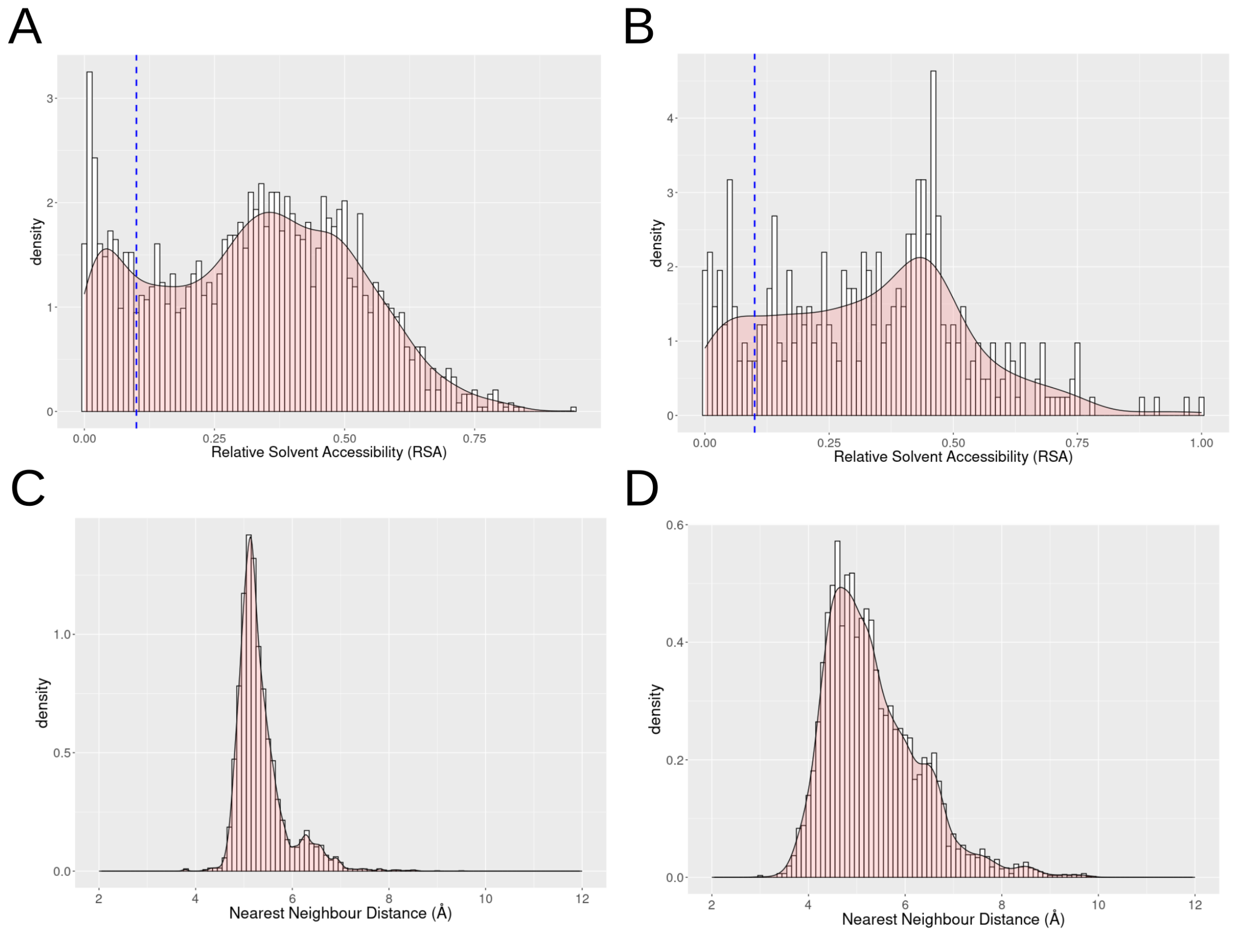


Figure S1: Distribution of Relative Solvent Accessibility (RSA) values for MHC-I (A) and MHC-II (B) residues at the interface with TCRs in TCR:pMHC structures. The vertical blue dashed line indicates an RSA of 0.1, corresponding to the 85th percentile in both MHC classes. Density plots were generated using a binwidth of 0.01. (C) Distribution of the nearest-neighbor distances between the Cα atom of solvent-exposed residues on the upward-facing pMHC surfaces from the set of pMHC-I and pMHC-II structures in TCR:pMHC complexes. Residues from the MHC β-sheet floor were excluded from this analysis: positions 1–50 and 85–136 for MHC-I, and 1–45 (α-chain) and 1–50 (β-chain) for MHC-II. (D) same as (C) but using side chain centroid. In both cases, the direct neighbors of the residues (i-1 and i+1) are not considered. Plots (C) and (D) consider an RSA cutoff of 0.1. Density plots were generated using a binwidth of 0.1.

To propose an appropriate threshold for determining edge formation between nodes in the graphs, we analyzed the distances between exposed residues on the surfaces of pMHC structures and their nearest neighboring residues. We observed an average distance of around 5Å when using side-chain centroids or Cα atoms (Supplementary Figure S1C and S1D). A distance threshold between 8.5Å and 10Å for forming edges between residue nodes would therefore be sufficient to fully cover the analyzed distances when using side-chain centroids or Cα atoms.

#### 2 Determination of chirality of node triads

To compose the descriptors of triads in tokens, a discrete class representing the chirality of the triad is determined. This descriptor assigns a pose-invariant (rigid-body invariant) and mirror-variant chirality label to each triad using local geometry derived from from Cα positions and side-chain direction vectors defined by Cβ coordinates (real Cβ when present, or a virtual Cβ for glycine).

For a triad of nodes $(v_i, v_j, v_k)$ with Cα coordinates $a_i, a_j, a_k \in \mathbb{R}^3$ and Cβ coordinates $b_i, b_j, b_k \in \mathbb{R}^3$, an unit normal vector of the Cα triangle is defined as the normalized cross product of two edges of the Cα triangle $\mathbf{n} = \frac{(\mathbf{a}_j - \mathbf{a}_i)\times(\mathbf{a}_k - \mathbf{a}_i)}{\|(\mathbf{a}_j - \mathbf{a}_i)\times(\mathbf{a}_k - \mathbf{a}_i)\|}$.

By using the Cβ coordinates, for each node $r \in \{v_i, v_j, v_k\}$ an unit side-chain direction is determined as $\mathbf{S}_r = \frac{\mathbf{b}_r - \mathbf{a}_r}{\|\mathbf{b}_r - \mathbf{a}_r\|}$.

Then, for each node, the signal is computed as $\sigma_r = \operatorname{sign}(\mathbf{n} \cdot \mathbf{S}_r)$. The majority signal across the three nodes is identified and when (majority_only = TRUE) (default), only nodes with the majority signal are kept. Exact zeros are mapped to +1 for numerical stability.

After majority filtering, let $K \subseteq \{v_i, v_j, v_k\}$ denote the set of residues whose side-chain directions agree with the majority side of the triad plane. The side-chain directions of the retained residues are combined by simple vector summation $\mathbf{v} = \sum_{r \in K} \mathbf{S}_r$.

The averaged side-chain direction is then defined as the unit-normalized vector $\mathbf{s} = \frac{\mathbf{v}}{\|\mathbf{v}\|}$. This normalization ensures that the subsequent chirality score depends only on the direction of the dominant side-chain orientation and is independent of the magnitude of side-chain agreement.

After computing the unit normal vector of the triad plane, $n$, and the averaged side-chain direction, $s$, we define a continuous chirality score as their dot product score $= \mathbf{n} \cdot \mathbf{s}$.

Because both vectors are normalized, the score lies in the interval $[-1, 1]$ and corresponds to the cosine of the angle between the triad plane normal and the dominant side-chain direction.

The final discrete chirality label is obtained as $\chi = \operatorname{sign}(\text{score})$ with exact zero values mapped to +1 by convention to ensure deterministic behavior.

To be able to perform the calculations when a glycine is presented in the node triad, a virtual Cβ is defined for the glycine residues, using only backbone atom coordinates, following a simple and stereochemically consistent geometric approximation. First, the the Cartesian coordinates of the backbone atoms N, Cα, and C are extracted and denoted as $r_n, r_{ca}, r_c$, respectively.

Two normalized backbone direction vectors are first computed as $\hat{\mathbf{n}} = \frac{\mathbf{r}_n - \mathbf{r}_{ca}}{\|\mathbf{r}_n - \mathbf{r}_{ca}\|}$ and $\hat{\mathbf{c}} = \frac{\mathbf{r}_c - \mathbf{r}_{ca}}{\|\mathbf{r}_c - \mathbf{r}_{ca}\|}$.

These vectors are combined to define an approximate side-chain direction $\hat{\mathbf{b}} = \frac{\hat{\mathbf{n}} + \hat{\mathbf{c}}}{\|\hat{\mathbf{n}} + \hat{\mathbf{c}}\|}$. The virtual Cβ position is then placed along this direction at a fixed distance from the Cα atom as $\mathbf{r}_{\text{virt}C\beta} = \mathbf{r}_{ca} - d_{ca\text{-}cb}\,\hat{\mathbf{b}}$, where $d_{ca} - d_{cb} = 1.522\text{Å}$ corresponds to the canonical Cα–Cβ bond length.

#### 3 MHCXGraph parameter description

| Parameter | Description | Type / Accepted Values |
|---|---|---|
| run_name | Define the name of the MHCXGraph run | string |
| run_mode | Define the mode of MHCXGraph execution | string (‘pairwise‘, ‘multiple‘, ‘screening‘) |
| dashboard | Activate this flag to generate an interactive dashboard for visualizing MHCXGraph results (recommended). | flag |
| reference_structure | Path to the reference structure when executing the screening mode | string |
| selectors | List of residues, chain or secondary structures to define the nodes in the protein surface graph | string |
| filter_triads_by_chain | Specify a chain to restrict triads to those having at least one node from the specified chain. This differs from providing a chain ID in selectors, as it allows focusing on a specific chain and its immediate surroundings (default = None). | string |
| max_gap_helix | Maximum number of residues between two helices to consider them as a single helix. This is useful when using secondary structure in selectors to treat long helical regions as continuous, even with small breaks (default = 0). | int |
| output_path | Path to output | string |
| max_chunks | Maximum number of input structures processed within a single group or batch during execution (default = 5). | int |
| debug_logs | Activate generation of logs for debugging purposes (default = false). | boolean |
| debug_tracking | Activate the generation of outputs from intermediate steps. Not recommended when running a large number of input proteins (default = false). | boolean |
| edge_threshold | Distance cutoff (Å) used to define edges between nodes. Larger values increase graph connectivity, generating more triads and increasing memory usage and computational time (default = 8.5). | float |
| node_granularity | Atomic representation used to compute 3D coordinates for each residue as nodes. | string (‘all_atoms‘, ‘ca_only‘, ‘backbone‘, ‘sidechain‘) |
| include_noncanonical_residues | Whether to consider non-canonical residues (e.g., modified amino acids) as nodes in the graph (default = true). | boolean |
| include_ligands | Whether to consider ligands as nodes in the graph (default = true). | boolean |
| include_waters | Whether to consider waters as nodes in the graph (default = true). | boolean |
| distance_bin_width | Width of the distance bins, used to define the number of classes for discretizing distance values. Higher values increase resolution but also the number of token classes (default = 2.0). | int |
| local_distance_diff_threshold | Maximum allowed difference between distances d1, d2, or d3 of a given triad compared to another triad to allow association (default = 1.0). | float |
| global_distance_diff_threshold | Maximum allowed distance difference between two non-adjacent node residues from an input protein compared to the corresponding associated nodes from another protein. This threshold is applied during the frame generation step (default = 2.0). | float |
| close_tolerance | Threshold value that defines the tolerance for distances d1, d2 or d3 to be placed at the center of the bin (default = 0.1). | float |
| triad_rsa | Whether RSA values are used in the triad token representation. If false, RSA is ignored and all triads share the same RSA class (default = false). | boolean |
| rsa_filter | Minimum RSA value required for a canonical residue to be considered exposed and included as a node. Zero means no filtering (default = 0.1). | float |
| asa_filter | Minimum ASA value required for a non-canonical residue, water or ligants, to be considered exposed and included as a node (default = 5). | float |
| rsa_bins_width | Width of the RSA bins, used to define the number of classes for discretizing RSA values. Higher values increase resolution but also the number of token classes (default = 0.3). | int |
| close_tolerance_rsa | Threshold value that defines the tolerance for RSA values to be placed at the center of the RSA bin (default = 0.01). | float |
| rsa_diff_threshold | Maximum allowed difference in RSAs of node residues from a given triad compared to another triad to allow association (default = 0.3). | float |
| watch_residues | Specify a protein residue to track during algorithm processing. Useful for debugging (default = None) | string |
| show_std_edges | Whether to display the standard deviation of distances between association nodes in the dashboard (default = False). | boolean |

Table 2: Summary of parameters and flags used for configuring MHCXGraph execution.

### 4 Example of Manifest file


```
{
  "settings": {
    "run_name": "test",
    "run_mode": "pairwise",
    "max_chunks": 5,
    "output_path": "/path/to/output",
    "debug_logs": false,
    "debug_tracking": false,
    "edge_threshold": 8.5,
    "node_granularity": "ca_only",
    "include_ligands": true,
    "include_noncanonical_residues": true,
    "include_waters": true,
    "triad_rsa": true,
    "rsa_filter": 0.1,
    "asa_filter": 5,
    "close_tolerance_rsa": 0.01,
    "local_distance_diff_threshold": 1.0,
    "global_distance_diff_threshold": 2.0,
    "distance_bin_width": 2,
    "close_tolerance": 0.1,
    "rsa_bin_width": 0.3,
    "rsa_diff_threshold": 0.3,
    "max_gap_helix": 5.0
  },
  "inputs": [
    {
      "path": "/path/to/input",
      "enable_tui": false,
      "extensions": [".pdb", ".cif"],
      "selectors": [
        { "name": "MHC1" }
      ]
    }
  ],
  "selectors": {
    "MHC1": {
      "chains": ["C"],
      "structures": {},
      "residues": {
        "A": [18,19,42,43,44,54,55,56,58,59,61,62,63,64,65,66,68,69,70,
        71,72,73,75,76,79,80,83,84,89,108,109,142,143,145,146,147,148,
        149,150,151,152,153,154,155,156,157,158,159,161,162,163,165,166,167,169,170,171]
      }
    },
     "MHC2": {
      "chains": ["C"],
      "residues": {
        "A": [31,51,52,53,55,56,58,59,60,62,63,65,66,67,69],
        "B": [56,57,59,60,61,62,63,65,66,67,68,69,70,71,72,73,74,77,78,81]
      }
    }
  }
}
```
